# Optical Coherence Tomography Reveals Self-Organizing Di-Fork Architecture of Mice Cutaneous Scars

**DOI:** 10.1101/181545

**Authors:** Biswajoy Ghosh, Mousumi Mandal, Pabitra Mitra, Jyotirmoy Chatterjee

## Abstract

Scientific studies report crucial impacts of biomechanical effectors to modulate wound healing either by scarring or regeneration. Further, the biological decision to predominantly favor the former is still cryptic. Real-time visualization of biomechanical manifestations *in situ* in scarring is hence necessary. Endorsed by nanostructural testing, synthetic phantom analysis, and computational simulations, we found strong mechanobiological correlates for Swept Source Optical Coherence Tomography (SS-OCT) speckles in mice cutaneous repair (full-thickness) up to 10 months. The theoretical basis of the optomechanics to provide insights into scar form-factor and evolution is proposed. Optomechanical changes have been considered as the resultant of intrinsic (e.g. fiber elastic modulus) and gross tissue mechanics (extracellular matrix (ECM)) in maturing scars. Non-invasive optomechanics supported with microscopic findings reveal scar’s cross-sectional self-organizing di-fork architecture. Dual-compartment heterogeneity of di-fork exhibits stress-evading features with a dichotomy in inhabitant cellular stress-fiber distributions. This differential interactivity of scar with adjoining tissues reflects its architectural intelligence to compensate tissue loss (hypodermis/muscle) by assembling into a di-fork. Gradual establishment of baseline shifted lasting mechanobiological steady-state, later in scarring, expose scar as an alternate stable state within the skin.

**Significance Statement:** Wound repair in mammals, predominantly culminates into function compromising scar that is occasionally fatal in vital organs. How the biological system often adopts scarring over a restorative regeneration is yet a conundrum. Wound and ambient mechanics play a pivotal role in deciding the healing fate. SS-OCT is hence demonstrated here as a non-invasive window to such mechanical manifestations during skin wound healing. This exposed gradual emergence of temporally maintained and stress-resilient di-fork architecture of the scar with differential neighborhood interfaces. Accommodation of such an alternate self-organizing steady-state of scar sheds light on its sustenance and paradoxical selection.

## Introduction

A scar is a result of inappropriate fibrillogenesis and fiber accumulation in mammals during wound repair. Alternatively, healing which completely restores original physiology is regeneration. Skin partial-thickness wounds heal regeneratively as opposed to life-long scarring in deeper assaults. Furthermore, scars in high mobility areas of the body (joints, abdomen, chest etc) are prone to aggressive fibro-proliferation e.g. hypertrophic scars, keloids etc [1]. Thus, wound milieu associated mechanics play a crucial role in determining healing fate. The central role of mechanical forces in regulating biochemical signals (mechanotransduction) in wound healing is well established [2]. But, causes of apparent immortality and “hyper” compensatory individualism of an emergent scar to mend resulting mechanophysiological imbalance needs investigation. The clinical load of fibro-proliferative disorders is not just restricted to skin but also several vital organs following repair of internal trauma e.g. heart attack, hepatic cirrhosis etc., and is also occasionally linked with cancer [3].

Insight into scar’s accommodation, despite its biological inappropriateness is revealed in the article emphasizing on wound-milieu mechanics. We investigated *in situ* scar development in murine skin during full-thickness excision wound healing with a Swept Source Optical Coherence Tomography (SS-OCT) for 10 months. Multiply-scattered photons documented modifications in tissue mechanical integrity during the healing course. This borrowed concepts from elastic Rayleigh/Mie scatters and inelastic stimulated Brillouin scatters (SBS) [4] resulting from laser radiation pressure (RP) [5]. Reflected dark field microscopy (RDFM) was adopted as a high resolution optical reflectance analogue to OCT. In-house phantoms with known properties also validated evaluated optomechanical connects. Atomic force microscopy (AFM) and nanoindentation (NI) experiments along with computational simulations assessed mechanical properties and stability parameters of scar. Our findings show that scars acquire a heterogeneity-modified and temporally-preserved stress resilient di-fork in cross-section (Fig. 1), to possibly acquire permanence in the system as an alternate stable state, suggestive of scar intelligence.

**Figure 1.**
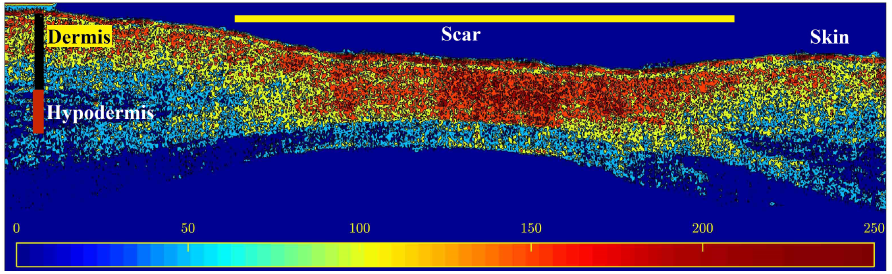
OCT intensity variations across scar di-fork.

## Materials and Methods

### SS OCT scan

Swiss albino mice (15, male, weight: 35-40 gm) were anesthetized (Ketamine/Xylazine) after chemical depilation of skin. Skin full thickness excision wounds (4-5 mm diameter) were created on either side of dorsum between 3rd and 6th Lumbar vertebrae. *In situ* wound tomograms were captured using SS-OCT with 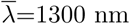, Δ*λ*=100 nm (OCS1300SS, Thorlabs-Inc., Newton, NJ, USA). Images were acquired every day post injury (dpi) up to 15 dpi and then at 30, 60, 120, 180, 240, and 300 dpi. Healing tissues collected at corresponding time-periods were formalin fixed for microscopy and mechanical analysis. The study was approved by Institute Animal Ethical Committee documented as IE-11/JC-SMST/1.14.

### Optical Microscopy

Hematoxylin/Eosin (HE) stained tissue sections were imaged under Leica DM750 (Leica Microsystems, Heerbrugg, Switzerland) at 10, 20 × objectives and Moticam1080 CMOS camera (Motic Asia, Hong Kong, China). Deparaffinised and dehydrated (DnD) tissue sections were imaged by stereo microscope (MVX10, Olympus, Tokyo, Japan) in Reflective dark-field (RDFM) mode at 1 × stereo objective and 2.0 zoom. For Immunohistochemistry, antigen retrieval of deparaffinised tissue sections was done in Tris-EDTA (pH 9.0) using EZ-Retriever V.2 (BioGenex, San Ramon, CA, USA) and immunostained with anti-*α*-SMA monoclonal antibody (Cat No.sc-32251, Santa Cruz Biotechnology, CA, USA) overnight at 4 °C. Alexa Fluor 594 conjugated secondary antibody (Cat No. A-11005, Molecular Probe, USA) was used for fluorescence immunodetection. Microphotographs were imaged with Zeiss Observer Z1 (Germany) and AxioCam, MRm camera (Carl Zeiss, Germany) at 10 and 20× objectives.

### Mechanical Imaging and Analysis

DnD tissue sections were imaged under Bruker Multimode 8 AFM system (Bruker Corporation, USA) visualized by PeakForce QNM (PF-QNM, Bruker Corporation, USA) mode. The nano-mechanical evaluation of tissue sections (DnD) was performed using the TI 950 TriboIndenter (Hysitron Inc., USA) with a diamond tip measured by loading of the tip up to 100 nm into the 4*µ*m sections followed by unloading. The reduced modulus (*E*_*r*_) and material hardness (H) were evaluated from the resultant load-displacement curve. Sample Young’s Modulus (*E*_*s*_) was calculated using 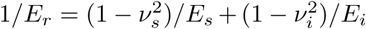, where *ν, E* are the Poisson’s ratio and elastic modulus of sample (s) and diamond indenter (i). *E*_*r*_ being the sample’s reduced modulus.

### Synthetic OCT phantoms

were prepared with agarose gel (0.2% w/v) and cellulose from Whatman filter paper grade1 (Sigma Aldrich, USA) after finely shredding, dispersing (shaking in water) and drying cellulose, eventually preparing a stock solution (0.8% w/v in water). Cellulose (0.2% v/v from stock) was boiled with agarose and 1 ml of mixture was casted into 12-well plates. Phantoms were air-dried.

### Image analysis

was done in MATLAB (R2016b, R2017b). GraphPad Prism 7 was used for plotting and statistical analysis (t-test, ANOVA). For OCT vs. time plot, OR (optical reflectance) was estimated from random patches (15 × 15) in maturing scars at specified time points. Patches selected from normal dermis adjacent to the scar, normalized reflectance alterations due to instrument fluctuations. OCT OR normalized to its corresponding dermis (*x*_*sc*_ − *µ*_*nom*_)*/µ*_*nom*_ was plotted against time. The baseline thus represented normal (y=0) at all time points. For OCT slope vs. intercept plot, signal change 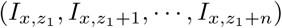 was plotted along scar depth *n* (A-scan amplitudes) to estimate OR changes. The depth-wise attenuation was equally divided into three zones from top to bottom. Linear regression fit was performed on each to be plotted as points in the slope-intercept plane. Texture and Histogram of oriented gradients were also done using inbuilt functions in MATLAB (details in SI Appendix)

### Computational Simulation

used COMSOL Multiphysics 5.1 platform. *Effect of RP on collagen fiber:* Cylindrical geometry represented collagen fiber (70 × 1000 nm) with E = 0.37 GPa [NI], density (*ρ*) = 1050 *kg/m*^3^, and *ν* = 0.4. The laser radiation pressure (F) was calculated using *F* = (2*P/c*)*r* cos *θ*, where *P* (power) = 10 *mW, c* the speed of light, *r* is reflectivity that ranges from 1 to 2 i.e. from completely absorbing to fully reflective material, *θ* is the angle of incidence (0 for normally incident laser) giving a maximum laser radiation force of 66.7 pN. *Effect of external forces on scar:* The scar di-fork as seen in tomogram was rendered in 3D (base diameter = 4mm) for a circularly healed scar (e.g. punch biopsy wounds). While the upper scar compartment (USC) assumed E=0.4 GPa (nanoindentation (NI)), *ρ*=500 kg/m3, *ν*=0.4, the lower compartment (LSC) allotted E=1 GPa (NI), *ρ*=1050 *kg/m*^3^, *ν*=0.4. Applied Force (0-300 N) in three different directions on the scar surface effected pressure, pull, and shear conditions.

## Results

### Optomechanics in OCT

The gray-intensity value of OCT speckles are estimates of optical reflectance (OR) backscattered from the corresponding tissue locations. Given the OCT resolution (9/12 *µ*m; air/water), OR illustrates micro-structural changes in scarring (Fig. 2a, SI Appendix Fig. S1). A broad range of elastic moduli was documented for both dermis and scar with nanoindentation experiment (Fig. 2e) with latter having an overall higher value. Similar observation was noted for OR of the two (Fig. 2f) plotted on determining speckle population fitted (Gaussian and Extreme-value models) model parameters i.e. mean (*µ*) and standard deviation (*σ*). Broader range of collagen fibril diameter (Fig. 2c,d) and orientation degree of freedom (Fig. 3c,d) was observed in intact dermis compared to scar under AFM and RDFM. To validate the optomechanical correlates, tissue mechanics was considered a cumulative function of (a) intrinsic mechanical properties of constituents (fibers e.g. collagen) and (b) overall matrix properties like hydration state and fiber density.

**Figure 2.**
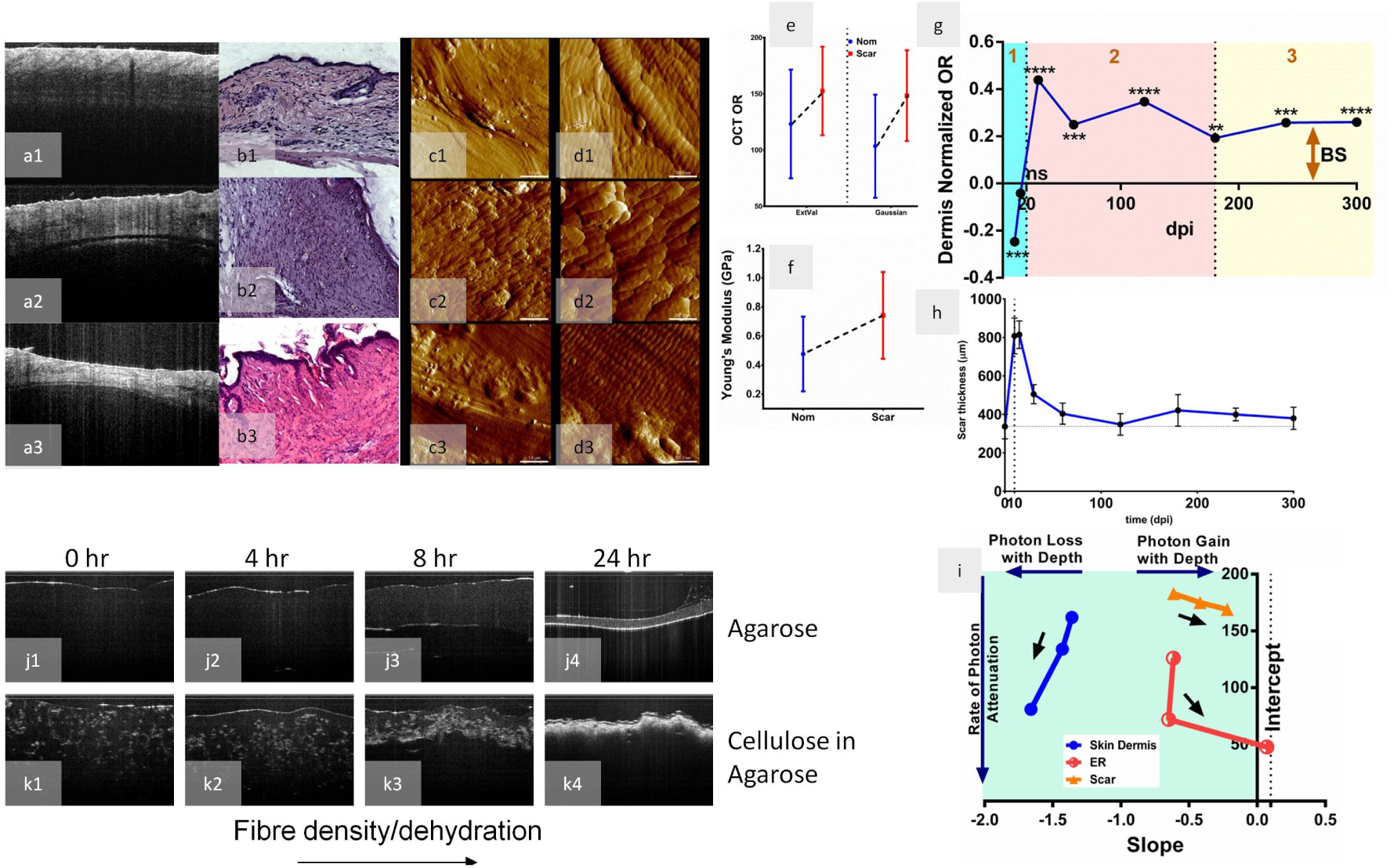
Illustration SS-OCT optical reflectance (OR) for non-invasive visualization of mechanical changes in scarring. **a** *in situ* mice OCT, **b** corresponding histology (HE stain), and AFM at **c** 10×10 *µ*m, **d** 2.5×2.5 *µ*m window size in **1** normal dermis **2** early remodeling (ER, 15dpi), **3** scar (180 dpi).**e** comparison of Young’s moduli (E) spread in skin dermis (N=30) and scar (N=30) from nanoindentation testing. **f** comparison the SS-OCT optical reflectance (OR) spread in skin dermis and scar (N>80k speckles × 5 ROI) assuming Gaussian and Extreme-Value population distributions. **g** temporal variations in OCT OR with scar maturation (N=30×8 time points) normalized at each point to adjacent normal dermis (baseline, y=0) with baseline shifts (BS) tested by two-tailed t-test (ns(P>0.05), *(P≤0.05), **(P≤0.01), ***(P≤0.001), ****(P<0.0001). **h** corresponding scar thickness (N=25) normalized to dermis thickness (origin). **i** slope-intercept plane locations of subsequent (back arrow) 1/3rd of the tissues from OCT depth scans representative of changes in depth-wise attenuation. The steepness of fall depict extent of attenuation and direction of shift (left/right) indicate increasing/decreasing OR attenuation with depth. **j, k** OCT of phantoms of agarose only and cellulose-agarose blend at progressive time points **1**,**2**,**3**,**4** of air-drying indicative of dehydration and simultaneous density (OCT intensity) gain. All error bars are *µ*±SD.

**Figure 3.**
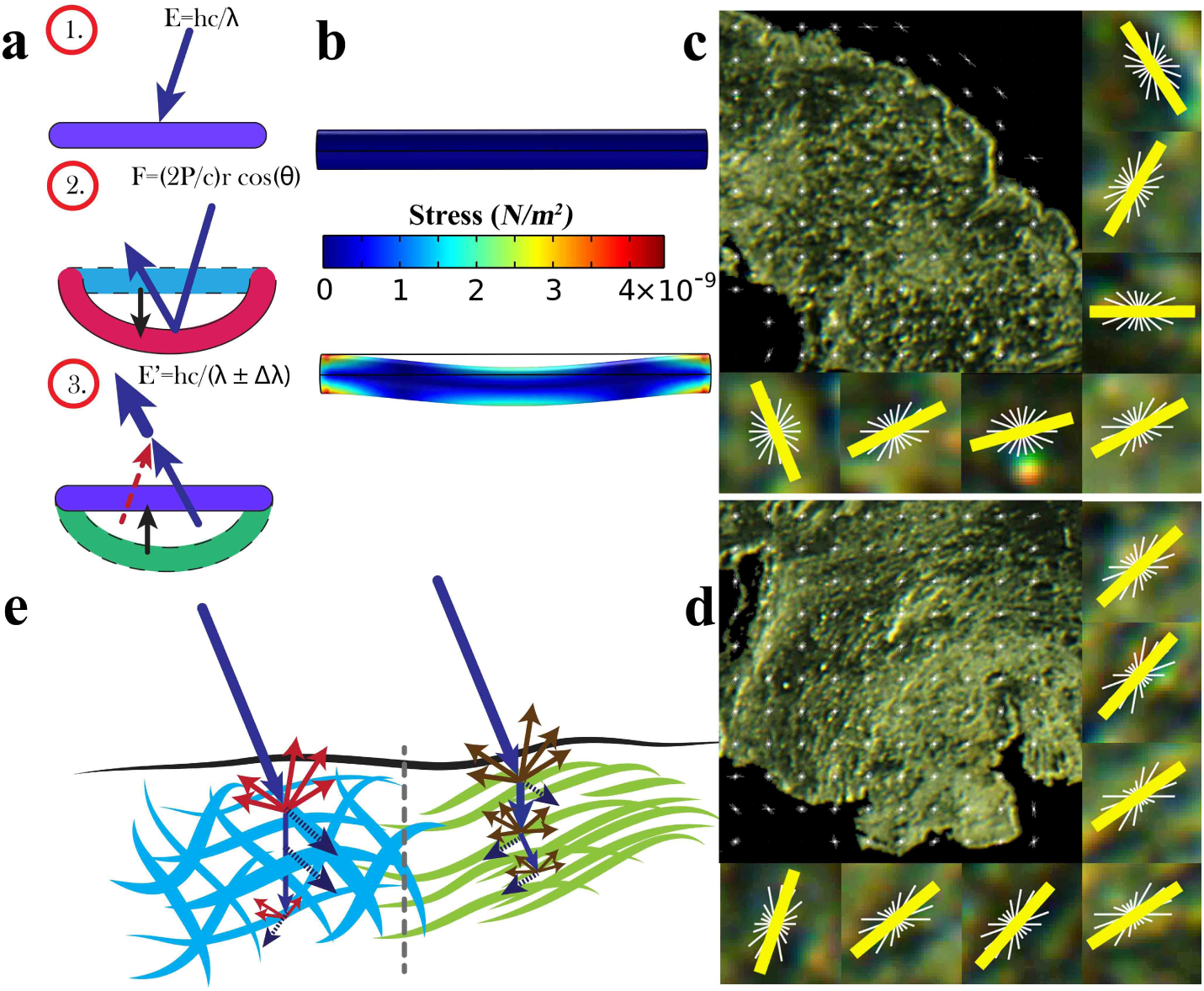
Schematic representation and theoretical correspondence of OCT OR *w.r.t.* both intrinsic and global tissue mechanics. **a-1**,**2** schematic of laser pressure induced fiber deformation and **a-3** the subsequent vibrations interfering with the photons altering its energy/wavelength to be reflected in OCT speckle intensities. **b** Simulation of the deformation in a 70×1000 nm fiber with E=0.37 GPa (top), laser force F=67 pN (bottom). **c** The fiber orientations calculated from RDFM images using HOG (rose plots) depicting tighter orientation in scar compared to normal dermis. **e** schematic illustration of differential photon interaction in normal dermis (left) and scar (right). Scar backscatters more photons owing to dense, parallel fibers with uniform diameter and low water content.

a. Endorsing the concept of radiation pressure, the SS-OCT with average NIR laser power of 10 mW, incident normally, theoretically exerts a force of up to 66.7 pN on the tissue components. Simulation illustrated effect of evaluated laser force on collagen fibril matched (shape, dimension and mechanical property) structure (Fig. 3a,b). An exaggerated skin mimicking phantom with fiber (cellulose, Young’s Modulus (E) ≈ 1-5 GPa [6]) embedded in water-retaining matrix (agarose gel, E ≈ 2 kPa-2 MPa [7,8]) illustrated proportionate OCT-OR response to the components at a given hydration state (Fig. 2j,k) i.e. cellulose displaying a higher OR than agarose even at completely dried state.
b. OCT was performed on phantoms of (i) only agarose and (ii) cellulose embedded in agarose (Fig. 2j,k) undergoing dehydration. The OR was recorded to increase with dehydration (and concentration/density) (Fig. 2j) as do their elastic moduli [7, 8]. Scar thickness was inversely proportional to OR during scarring (Fig. 2g,h).

### Spatio-temporal variations in scar optical reflectance

The OR measures were determined along (a) healing tissue depth and (b) temporal axis of wound repair up to 300 days post injury (dpi) (SI Appendix Fig. S1).

a. OCT OR change with depth (predominantly attenuation) in tissues (dermis, ER, scar) was plotted and fitted to a line (SI Appendix Fig. S2). To get a better resolution of signal change with depth, a slope vs. photon attenuation for every progressing one-third depth of tissue was plotted (Fig. 2i). Normal dermis displayed progressive attenuation (left shifted i.e. increasingly negative slope) with significant signal loss between steps (high vertical distance between progressive marks) while the scar progressed towards the right with negligible vertical shift between points implying much lowered signal attenuation. The first 2/3rd ofearly remodeling (ER) tissue displayed high attenuation (left shift) contrary to the right shift in its lowest zone. These indicate the gradual emergence of a high OR zone in deeper scar region resulting in lowered attenuation of scar.
b. Post injury the wound bed OR was found to be significantly low (Fig 2g). At 15 dpi, the OR nearly matched normal followed by a spiked rise within the next 15 days. Fluctuations (rate change) in OR gradually declined with time and reached a plateau (180 dpi onwards) that was distinctly shifted from the normal dermis baseline.

### Scar’s modified heterogeneity

RDFM of dehydrated tissues sections (4 *µ*m) illustrated dual compartment within scar. Distinct upper (USC) and lower scar compartments (LSC) with higher reflectance (Fig. 4e), texture (range, entropy) (Fig. 4f,g) and Young’s modulus (Fig. 4d) in the latter. Healing actuators i.e. the Myofibroblasts (MFB) were found to express even in matured scars (180 dpi) as seen by marker *α*-SMA immunofluorescence (Fig. 4a). Further, a clear morphological difference in *α*-SMA expression of MFBs was seen in scar USC and LSC (Fig. 4b,c).

**Figure 4.**
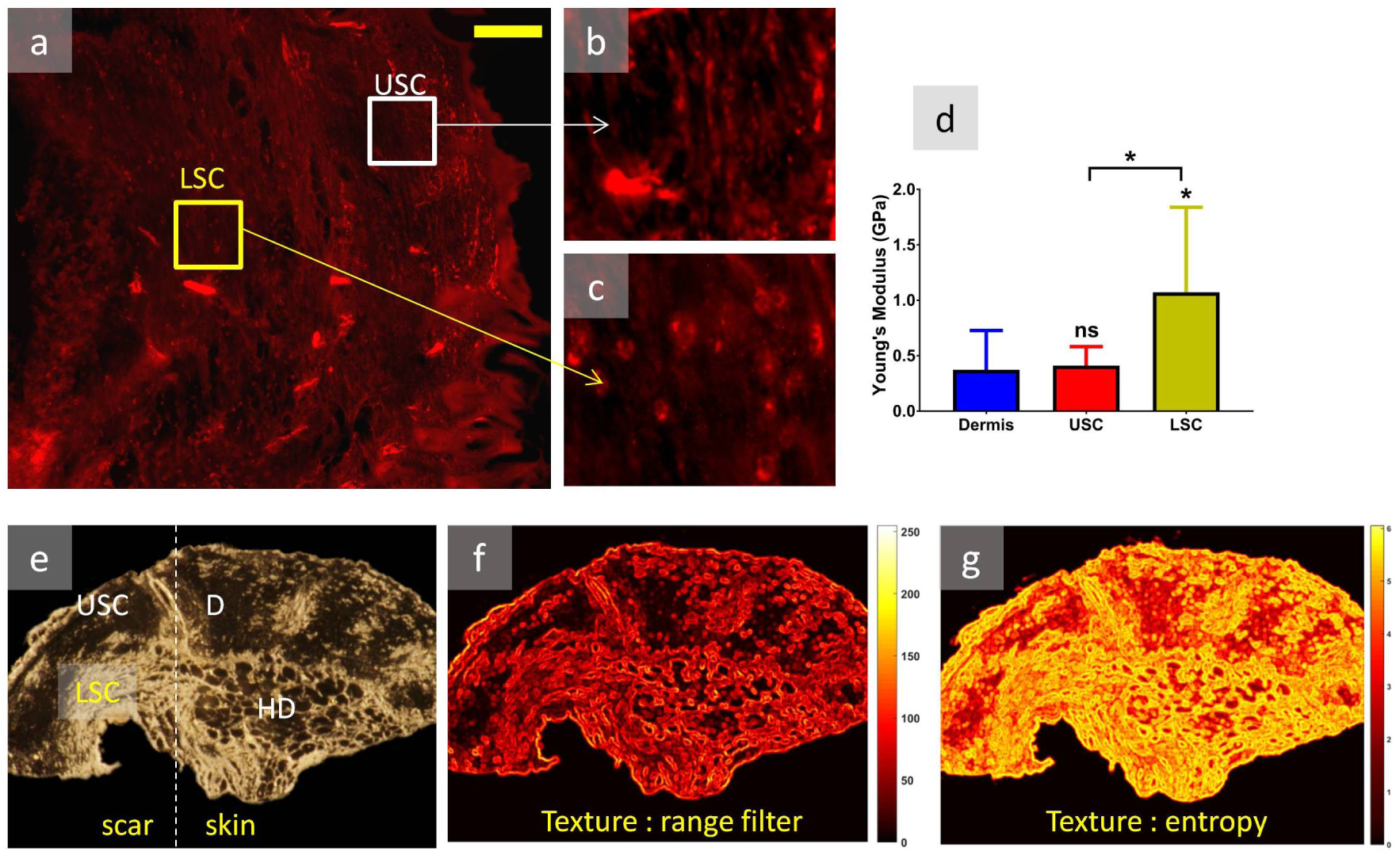
Mechanical dichotomy in scar’s modified compartmentalized heterogeneity and its impact on cells. **a** expression of *α*-SMA in scar. **b** *α*-SMA positive MFBs in USC predominantly demonstrate spindle shaped appearance, **c** circularly spread distribution of *α*-SMA in the LSC. **d** nanoindentation results display 2-fold increase in Young’s modulus (E) in LSC compared to USC and normal dermis (N= 10 per group) **e** RDFM image of the scar-skin junction section demonstrating a higher reflectance from LSC. **F**. **f**,**g** textural map of the scar region based on overall intensity range and entropy respectively highlighting differences in the scar compartments for RDFM image.

### Scar di-fork architecture

The optomechanical nature of OCT signals for scars were visualized as a heat map of its OR measures (Fig. 1). The 4-10 month old scar OR were prominently found extending into the intact dermis above and muscle layer below leaving a cleft in the middle constituting mainly of neighboring hypodermis (Fig. 5a). The Hypodermis was absent in scar. The di-fork architecture was simulated in 3D as illustrated in Fig. 5b,c. The emergent structure was subjected to shear, pressure and pull (Fig. 5d,e,f). In all cases the structure recorded deformation, dissipating majority of the stress in the neighboring fat-rich hypodermis attributed to the di-fork form.

**Figure 5.**
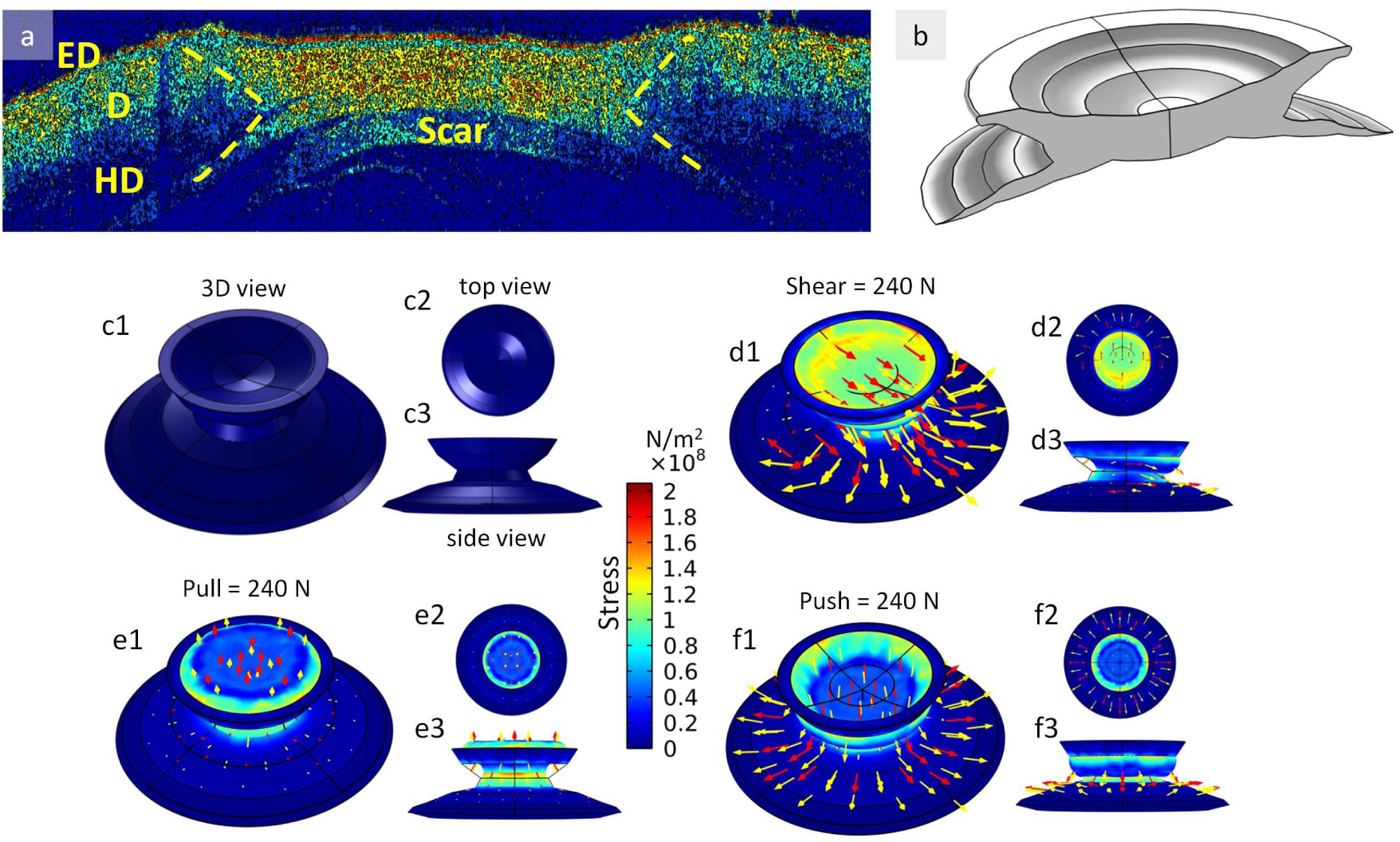
The di-fork architecture of scar and its stress distribution features. **a** Scar OCT heat-map displaying forking arms stretching both above and below the adjacent hypodermis (ED-epidermis, D-dermis, HD=hypodermis). **e** simplified 3-D representation of the scar with di-fork in cross section. **c-1**,**2**,**3** are the 3D, top and side view of a simulated circularly healed scar with differential mechanical properties in USC and LSC (4mm base diameter). **d**,**e**,**f** simulates stress distribution and deformation of scar on impacts of shear, pull(vertical) and push (vertical) forces at F=240 N using evaluated mechanical properties.

## Discussions

This article highlights maturing cutaneous scar architecture and proposes SS-OCT as a tool for extracting some of its physiologically relevant facets *in situ* using optomechanics. OCT optomechanics here is the optical projection of cumulative tissue mechanics, and not absolute mechanical properties. This paper tracks maturation of scar architecture as a “system within a system” while illustrating relevant insights on its permanence.

### Optical scattering and tissue mechanics

The SS-OCT uses NIR laser (1300 nm) lying in the second IR window for optimum penetration in biological specimen [9]. While the backscattered photons contribute to the image intensity, tissue-specific photon attenuation reduces it. The detailed skin-photon interaction optics is beyond the scope of this article [suggested read [10]]. Lasers potentially exert radiation pressure (RP) and are used in maneuvering small particles e.g. optical tweezers [11] and depends on laser power (P), reflectivity of the interacting material (r) and the angle of incidence (*θ*) [12]. Our simulations substantiate effect of radiation pressure of the OCT laser on nano-scale skin collagen fibrils (Fig. 3b). The backscattered photons after tissue-interaction are received with their residual energy on the SS-OCT photo-detectors where they are converted to equivalent electrons for image formation. Since OCT images multiply-scattered photons [13], it paints a deterministic picture of the source of spatial intensity distributions unlike random singly-scattered phenomenon. The mechanical property of ECM depends on both material elastic modulus and density. High density scattering components correspond to high probability of photon backscatters (Rayleigh, Mie) reaching the detector, represented in image as higher intensities in respective regions (Fig. 3e). Furthermore, materials undergo elastic moduli dependent deformations due to laser radiation pressure resulting in vibrations (phonons) that interfere with the light i.e. inelastic stimulated Brillouin scattering (SBS), altering its wavelength/energy (Fig. 3a,b) and thereby modulating image intensity. The photon-phonon interference has been directly used to estimate mechanical properties of objects including collagen fibers [14] and improve OCT resolution [13]. In fact, optical coherence elastography (OCE) measures differences in tissue stiffness by external perturbations (loading/unloading, ultrasound etc.) followed by speckle tracking [15]. In our study, the vibrations are inherently caused by SS-OCT laser RP [12] inducing matrix deformation and subsequent photon-phonon interfered SBS [4] being reflected as intensities on the SS-OCT image. In-house phantoms designed to represent skin with fiber (cellulose) in a water retaining matrix (agarose gel) show OCT image intensities (Fig. 2j) to proportionately increase with elastic moduli. As material density/volume fraction (inverse of porosity) also effects elastic modulus positively [16], higher density of cellulose fibers display pronounced OR intensities (Fig. 2k). Thus, higher scar OR can be attributed to either or both of increased *E*_*collagen*_ and fiber density.

### Optomechanics of tissue hydration

Fiber density can be caused by rapid polymerization (relative to degradation) or water loss. While the proteoglycans (PG) form the water retaining ground matrix for cushioning cells and fibers, extensive water bridges are bound to collagen fibers [17] with structural and functional relevance [18]. Thus, water content also dictates tissue mechanics. Vast fluctuations in elastic moduli (≈ 10^6^*Pa*) have been reported within same materials (including collagen) based on their hydration state [7, 8, 14]. Water absorbs most SS-OCT NIR energy (by vibration in O-H stretching), mechanically resembling energy loss by photon-water interaction (absorption). Low reflectivity (r) in water-rich materials, in terms of RP, renders low photon energy reaching the detector, effectively lowering OR intensity. Agarose gel-only phantoms display increasing OR with dehydration (increasing E) (Fig 2j). Further, in fiber embedded gel phantoms (AgCel), the effect is even more pronounced due to effective gain in fiber density with water loss. Early healing (ER) phase (granulation tissue) is full of high moisture containing hyaluronic acid (HA) [19] thus display a low OR in OCT (Fig. 2a2,g). Initial studies have reported a mirror symmetry (inverse relation) between HA content and mechanical strength [20] in healing tissues. The OR peak (15 dpi onward) observes an inverse correspondence to shrinkage of scar thickness (Fig. 2g,2h) suggesting water depletion. This is due to simultaneous reduction in HA (replaced by polymerization boosting sulfated proteoglycans [21], decreasing polymerization regulating decorin [1]) and increased degradation resistant hydroxyl-lysine cross-linking [22] (shrinking tissue by exuding-out fluids) in parallel collagen bundles during scarring.

### Di-fork architecture heterogeneity

AFM (Fig. 2d1,d3) alongside RDFM (Fig. 3c,d) images illustrate the reticular and parallel arrangement of collagen in dermis and scar. Furthermore, fibril diameter in scar is observed to be uniform compared to normal dermis (Fig. 2d1,d3). Scar’s geometric evenness and tighter angular orientation may be contributory to uniformity in OCT scans (Fig. 1) attributable to homogeny in elastic light scattering. Although scars appear heterogeneity compromised, OCT slope-intercept plot points out the gradual enhancement in photon gain (right shift) (Fig. 2i) with scar depth (SI Appendix Fig. S2) as completely opposed to dermis with the usual increasing attenuation. Further, the scar (tomogram) extends both above and below the adjacent hypodermis (Fig. 5a) having a di-fork appearance with upper scar compartment (USC) merging into neighboring dermis and lower (LSC) clutching onto muscle layer. RDFM yields OCT-homologous reflected photons from structures in scar’s sectional view demarcating the compartments clearly (SI Appendix Fig. S3, Table S1). Texture (Fig. 4f,g) and intensity distributions (Fig. 4e) in both compartments depict USC and LSC having uniformly low and high reflectance while dermis spans the entire range (more intra-tissue variations). Mechanically, *E*_*USC*_ is closer to *E*_*dermis*_ while being significantly deviated from *E*_*LSC*_ (Fig. 4d). This establishes the premise of scar’s property of dual biomechanical compartmentalization depth-wise to possibly achieve effective coordination with interacting normal tissues towards mechanically compensating their absence.

### Di-fork mechanobiology

Fibroblast to myofibroblast (MFB) differentiation (manifested by stress fiber *α*-SMA expression) besides, inflammatory signals, is induced by mechanical forces [23] and helps in wound contraction by laying collagen. As acute inflammatory signals diminish after ECM restoration (15-20 dpi), MFBs die by apoptotic regulation [3]. However the prolonged cellular *α*-SMA expression (180 dpi) suggests mechanical stress-induced MFB production of the scar. Furthermore, *α*-SMA morphologies in USC and LSC are indicative of stress fiber response to diverse mechanical stresses i.e. spread-out in higher mechanical stress and narrow/compact in lower [24]. The spread out (ellipsoidal) *α*-SMA expression in LSCs and fusiform (narrow, tailed) ones in USCs (Fig. 4a,b,c) advocate differential cellular effects in the di-fork compartments. This is analogous to the differential sub-populations of fibroblast morphologies localized in dermal proximities (papillary, reticular, follicular) [25]. Literatures further indicate apoptotic regulation of MFBs on releasing mechanical stress. A possible explanation to the prolonged *α*-SMA expression could be that the matrix-stress triggered MFBs which being potent cytokine attractors [3] help maintain inflammatory signals (chronicity) adding to further MFB differentiation. The collective pool of MFBs (induced mechanically [26] and by inflammation [3]) lays out more scar collagen thus maintaining scar mechanics and entering a closed loop vicious cycle.

### Self-organization of di-fork attaining mechanically conducive alternate steady state

The di-fork, in perspective resembles self-organization wherein a relatively chaotic event (wound) assembles into a definite form using feedback from exploratory local internal fluctuations to gradually attain a dynamic equilibrium. The full thickness wound after ER (15dpi) undergoes rapid collagen polymerization and water loss (up to 30 dpi depicted as OR spike Fig. 2g) inherently. Thereafter without any known exception di-fork architecture emerges which enters into a phase of fluctuating matrix modification (30 dpi-120 dpi OR fluctuations, Fig. 2g) effected by plausible biological feedbacks from scars’ mechanical alteration (Fig. 4d). The di-fork dissipates majority of stress towards the adjoining fat-rich hypodermis (Fig. 5d,e,f). In the context of achieving permanence in cutaneous environment, scar as an emerging system has to iteratively nurture itself (fiber assembly, composition, compartmentalization) using feedback loops and addresses loss of crucial structures (hypodermis and muscle layers) with forked geometry and mechanobiological dichotomy. Thus it improvises effective tissue interfacing reflected as temporally maintained di-fork’s steady state (180 dpi onwards, Fig. 2g) perhaps on realization of an optimized mechanical condition.

## Conclusion

The inherent relevance of SS-OCT to document strong mechanical correlates in wound healing as the cutaneous scar matures in the living system instrumented investigation into the perpetual aspect of scarring. Scar, although being broadly classified into types essentially exists in a complex continuum of structure and time, understanding of which may effect personalized wound-care. The scar di-fork pronounce consistent forces in the wound milieu and optimizes an architecture to maintain the mechanical balance with the newly deposited ECM and lack of buffers like subcutaneous fat. The unusual mechano-stable features of the architecture can be a potential cause of scar’s longevity. Reverse engineering applications of the structure in mechano-intervention and tissue engineering may draw crucial inputs in restoration of normal skin architecture (scar-less healing) i.e. more regenerative. As the early healing stages do not illustrate this di-fork when ECM is freshly laid, it sets the quest for a temporal switch after which healing progresses to an irreversible and everlasting scar.

## Acknowledgments

The study was funded jointly by Indian Institute of Technology, Kharagpur (IIT/SRIC/SMST/CIT/201 15/122, dated 24.07.2014) and Department of Biotechnology, Government of India (BT/PR7961/MED/ 32/280/2013, dated: 18.06.2014). BG was funded by Ministry of Human Resource Development, Government of India (F. NO. 4-23/2014 – TS.I, dated: 14.02.2014).

## Notes

### Competing Interest Statement

The authors have declared no competing interest.

### Summary of Updates

The version of the manuscript makes few revisions in figures Fig 2, 4, 5 for a better understanding of the paper. Few minor text corrections have been made to reduce complications. A minor change in the title of the manuscript.

