## Supplementary material for "Optical Coherence Tomography Reveals Self-Organizing Di-Fork Architecture of Mice Cutaneous Scars": SI Appendix

### Supporting Information Appendix

#### Image Analysis Methods for RDFM images

##### Texture Analysis

***Range Texture:*** For an input image $I$ of size $m \times n$, the range filtering operates on every image pixel $I_{x,y}$ such that every output pixel $O_{x,y}$ gives a range value i.e. $I_{max} - I_{min}$ in the $3 \times 3$ pixel neighborhood of the input pixel $I_{x,y}$. The gray-scaled RDFM image was subject to range filtering operation to visualize the spatial texture with an appropriate color map displaying values spanning from 0 to 255. When a $3 \times 3$ neighbor of an input pixel all have the same values, it will give a 0 range, while in the event of the neighborhood to contain both brightest (255, high reflectance) and darkest pixels (no reflectance), the operation gives a range of 255.

***Entropy Texture:*** Entropy is the measure of randomness, in this context, for the image. For an input image $I$ of size $m \times n$, entropy filtering operates on every image pixel $I_{x,y}$ such that every output pixel $O_{x,y}$ gives the entropy value i.e. $\sum(p \log_2 p)$ for a $9 \times 9$ pixel neighborhood of that input pixel $I_{x,y}$. Here, $p$ is the probability of a gray intensity value (0 to 255) within the neighborhood determined from its histogram (probability density function). Thus, a neighborhood with more number of intensity bins (higher contrast) will have a higher value of randomness or entropy. The gray-scaled RDFM images were operated by entropy filtering to determine regions of higher and lower contrast of reflectance.

##### Histogram of Oriented Gradients:

This is a widely used feature descriptor algorithm in computer vision to visualize angular distribution of gradients. The algorithm is employed here to decipher the angular variation of collagen fibers in RDFM images. A region of interest (ROI) was selected ($\approx 770 \times 770$ pixels) from the main image. The ROI image was divided in cells (equally spaced compartments) of size $52 \times 52$ such that each cells can capture sufficiently resolvable gradient changes. The algorithm uses a 1-D sobel operator to evaluate the horizontal ($G_x$) and vertical gradients ($G_y$). Two $52 \times 52$ matrix are generated for gradient magnitude, $G = \sqrt{g_x^2 + g_y^2}$ and gradient direction, $\Theta = \arctan(g_y/g_x)$, where $g_x$ and $g_y$ are the pixel values of the horizontal ($G_x$) and vertical gradient ($G_y$) matrices. Nine angular bins are assigned 0, 20, 40, 60, 80, 100, 120, 140, 160 and magnitude values corresponding to the angles are filled in the appropriate angular bins. For gradient values with angles between two successive bins, are proportionately distributed. The evaluated histogram is visualized as a rose plot (polar representation) with complementary angles assigned to the same height value (0-180, 90-270). The dominant angle of the rose plots was highlighted.

### Supporting Figures and Tables

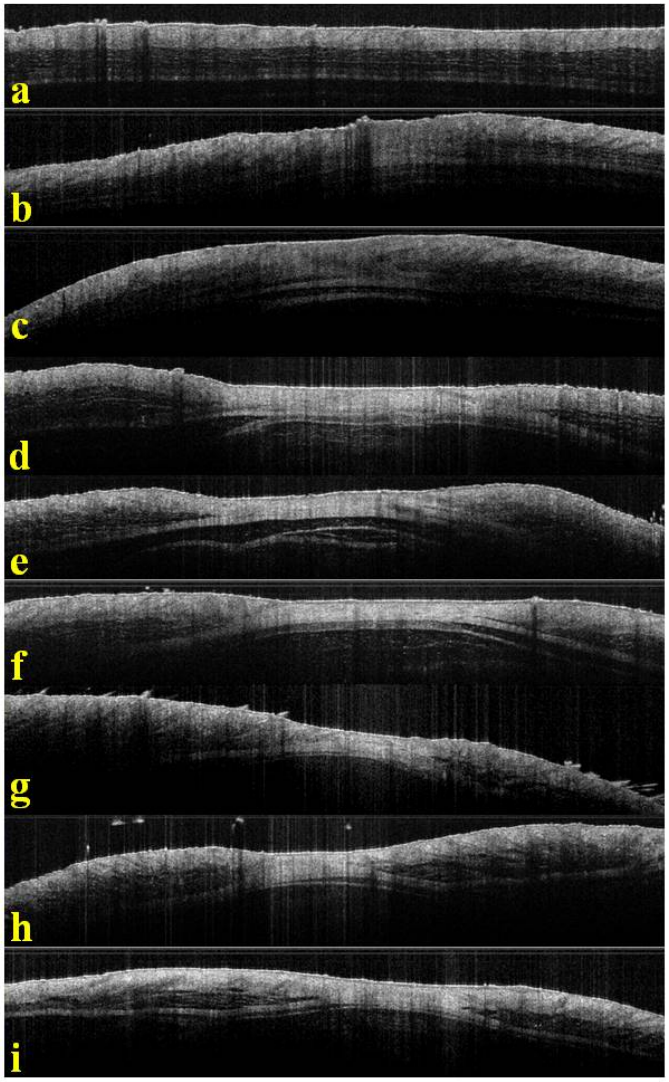


Figure S1. OCT scans (not spatially matched) of (a) normal skin and wound bed in different healing stages at time points as days post injury (b) 10 dpi (c) 15 dpi (d) 30 dpi (e) 60 dpi (f) 120 dpi (g) 180 dpi (h) 240 dpi (i) 300 dpi demonstrating the modified OR and prominence of the di-fork.

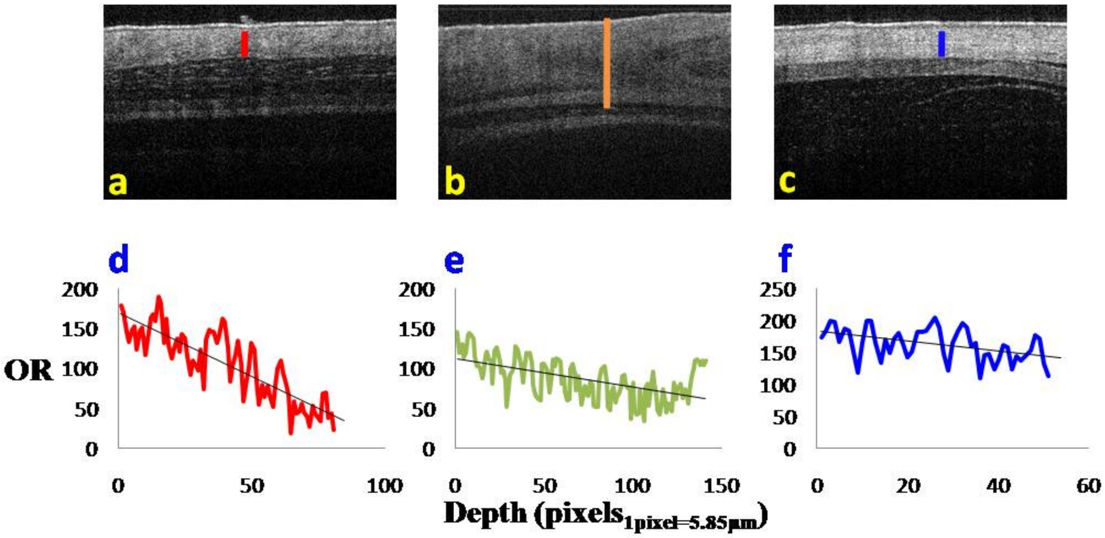


Figure S2. The figure displays the optical attenuation with tissue depth for (a) normal dermis (b) early remodeled scar of 15 dpi (c) matured scar of 180 dpi. The normal shows a (d) uniform attenuation while for ER(e)attenuation is followed by gain, and for scar (f) the attenuation almost balances the gain represented as a constant trend line.

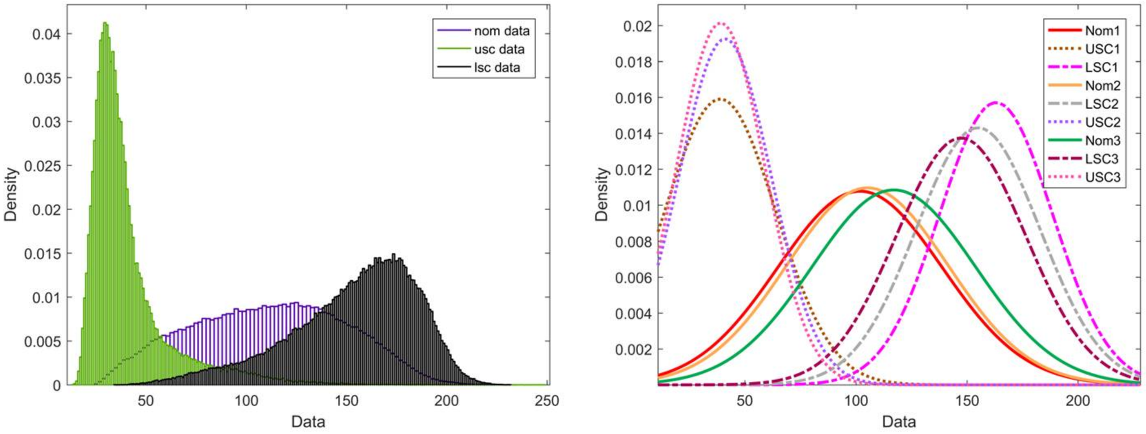

Figure S3. The distribution of reflected photon intensity in RDFM (N>50k). *left* shows the normal dermis (blue) distribution is spread across the entire intensity spectra while the USC (green) and LSC (black) have a more polarized distributions as an indication of respective heterogeneity spread. *right* are the Gaussian envelopes of the distributions of normal dermis, USC, and LSC.

**Table S1.** Tabulation of the mean and standard deviations of the four populations (N>50k). The normal has the largest intra-tissue variation ($\sigma$) followed by LSC and USC.

| | **Normal Dermis** | | **USC** | | **LSC** | |
|---|---|---|---|---|---|---|
| | $\mu$ | $\sigma$ | $\mu$ | $\sigma$ | $\mu$ | $\sigma$ |
| **Population 1** | 39.08 | 19.78 | 163.09 | 25.39 | 117.035 | 36.76 |
| **Population 2** | 41.06 | 20.71 | 154.99 | 27.87 | 105.142 | 36.37 |
| **Population 3** | 38.88 | 25.06 | 147.32 | 29.03 | 101.824 | 37 |
| **Population 4** | 40.72 | 20.94 | 152.01 | 31.64 | 110.33 | 38.085 |
